# A wound-induced differentiation trajectory for neurons

**DOI:** 10.1101/2023.05.10.540286

**Authors:** Ryan E. Hulett, Andrew R. Gehrke, Annika Gompers, Carlos Rivera-López, Mansi Srivastava

## Abstract

Animals capable of whole-body regeneration can replace any missing cell type and regenerate fully-functional new organs, *de novo*. The regeneration of a new brain requires the formation of diverse neuronal cell types and their assembly into an organized structure and correctly-wired circuits. Recent work in various regenerative animals has revealed transcriptional programs required for the differentiation of distinct neuronal subpopulations, however how these transcriptional programs are initiated upon amputation remains unknown. Here, we focused on the highly regenerative acoel worm, *Hofstenia miamia*, to study wound-induced transcriptional regulatory events that lead to the production of neurons. Footprinting analysis using chromatin accessibility data on an improved genome assembly revealed that binding sites for the NFY transcription factor complex were significantly bound during regeneration, showing a dynamic increase in binding within one hour upon amputation specifically in tail fragments, which will regenerate a new brain. Strikingly, NFY targets were highly enriched for genes with neuronal functional. Single-cell transcriptome analysis combined with functional studies identified *sox4^+^*stem cells as the likely progenitor population for multiple neuronal subtypes. Further, we found that wound-induced *sox4* expression is likely under direct transcriptional control by NFY, uncovering a mechanism for how early wound-induced binding of a transcriptional regulator results in the initiation of a neuronal differentiation pathway.

**Highlights:**

- A new chromosome-scale assembly for *Hofstenia* enables comprehensive analysis of transcription factor binding during regeneration
- NFY motifs become dynamically bound by 1hpa in regenerating tail fragments, particularly in the loci of neural genes
- A *sox4*^+^ neural-specialized stem cell is identified using scRNA-seq
- *sox4* is wound-induced and required for differentiation of multiple neural cell types
- NFY regulates wound-induced expression of *sox4* during regeneration

## Introduction

Animals capable of whole-body regeneration can replace all missing cell types, including those within the nervous system^1,2^. These species present an opportunity to identify the molecular and cellular mechanisms of neural specification and differentiation that enable regeneration of entire new brains in adult animals. While these processes have been previously studied during embryonic development across diverse metazoan systems^3–14^, the transcriptional regulatory decision-making connecting the early steps of regeneration with neural specification and subsequent differentiation is not well understood. Here, we sought to identify changes in chromatin accessibility and transcription factor binding that lead to the formation of neurons during whole-body regeneration.

Across animals, many lineages include species capable of whole-body regeneration and extensive adult neural regeneration (Fig. 1a). Non bilaterians, such as cnidarians^15–22^, and ctenophores^1,23–27^, which lack centralized nervous systems, are well-known for their ability to regenerate nervous systems. Among bilaterians, which have centralized nervous systems with brains, classic examples of animals with robust neural regeneration include enchinoderms^28–30^ and hemichordates^31–33^- comprising the group of animals known as ambulacrarians, as well as planarians^34–42^. Extensive work has been done in planarians to understand the genetic requirements for neural regeneration and putative specialization of neural-primed stem cells^36,40,42–50^. In the highly regenerative cnidarians *Hydra* and *Nematotella,* recent work provided a comprehensive transcriptional characterization of the adult nervous system including differentiation pathways, inferring the molecular trajectories for the formation of differentiated neurons during homeostasis^51,52^. However, we do not know how programs mediating neuronal regeneration are activated in these systems.

**Figure 1.**
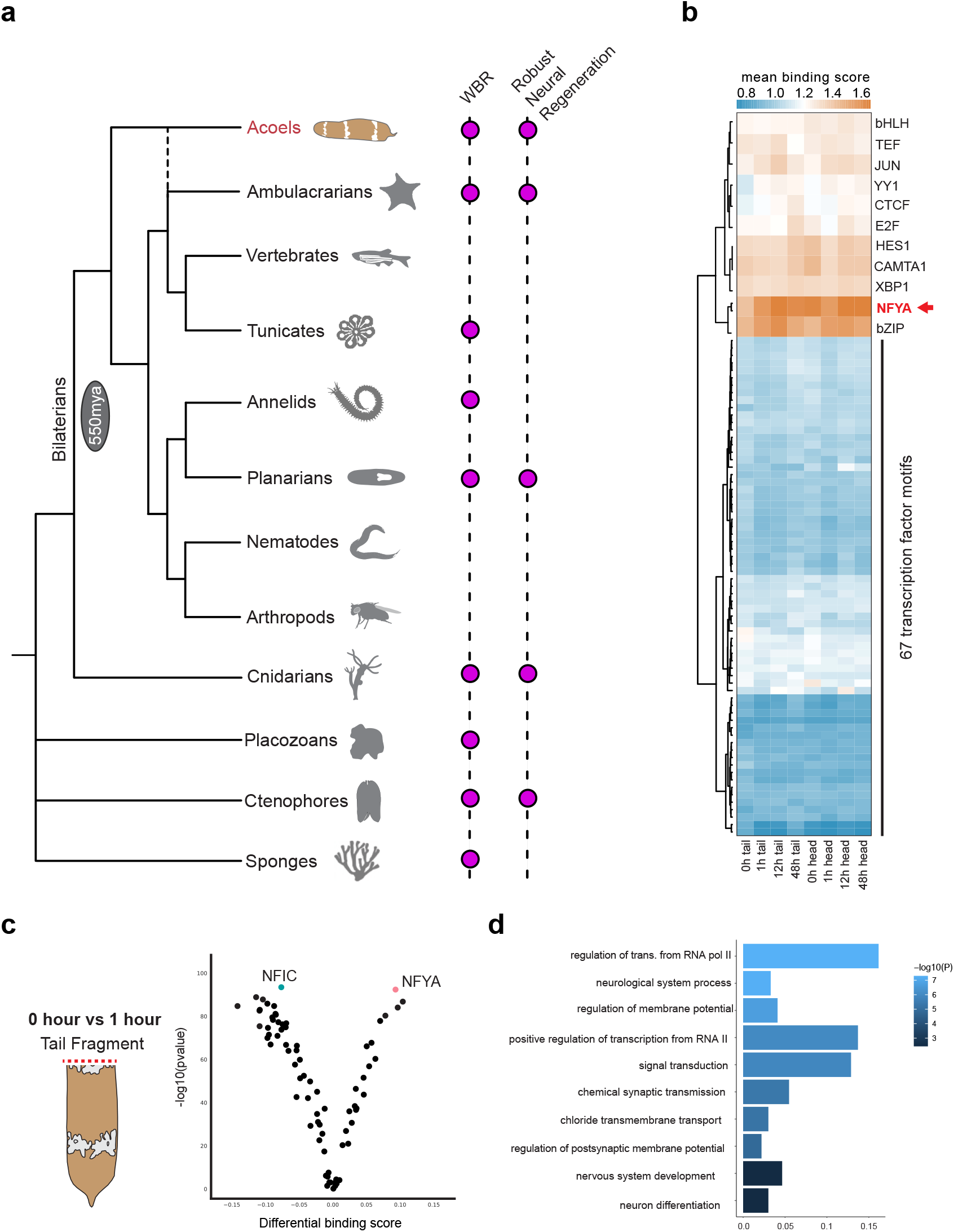
NFY is the most significantly bound motif in the *Hofstenia* genome during regeneration. (A) Schematic phylogeny of select metazoan lineages based on published literature^108^ identifying animals capable of whole-body regeneration (WBR) and robust neural regeneration. Labeled with the origin the Bilateria ∼550 million years ago (mya). (B) Heatmap showing the mean binding score calculated at the binding sites associated with the curated list of 78 *Hofstenia* transcription factors. In head and tail fragments sampled during regeneration, NFY is the most differentially bound transcription factor binding site in the *Hofstenia* genome. (C) Volcano plot showing that NFIC is the most bound motif at 0hpa and NFYA is the most bound motif at 1hpa in the regenerating tail fragment in the *Hofstenia* genome. (D) Dynamically bound NFY sites at 1hpa compared to 0hpa are linked to genes associated with neural function. Gene ratio is the number of genes in a particular biological process category over the total list of genes.

Acoels represent a bilaterian lineage with diverse neural architectures, are also capable of extensive neural regeneration^53–55^, and hold a phylogenetically informative position, as a sister group to all other bilaterians^56–64^ or ambulacrarians^63,65–68^. Studies of regeneration in acoels can provide insight into mechanisms of neural specification and differentiation during whole-body regeneration as well as their evolution. The acoel, *Hofstenia miamia*, regenerates its brain and nerve net^53^ and is amenable to mechanistic studies of regeneration^64,69^. Chromatin accessibility data have also enabled the identification of gene regulatory networks for regeneration in this system^55,70^. Therefore, we sought to understand the regulatory decision-making for neuronal regeneration in *Hofstenia*.

Here, we generated a chromosome-scale genome assembly and undertook a comprehensive study of transcription factor binding during regeneration. This showed that the NFY transcription factor complex becomes dynamically bound within an hour upon amputation genome-wide, particularly at the loci of nervous system-associated genes. Further, functional studies showed that NFY is required for brain regeneration. Next, we conducted a systematic analysis of single-cell transcriptome data focusing on neuronal cells and identified *sox4* as a wound-induced factor that marks neuronal progenitors and is required for the formation of diverse neuronal cell types. Finally, we found that NFY is likely a direct transcriptional regulator of wound-induced *sox4* expression. Altogether, this work identifies a path from a wound-induced change in chromatin binding to specification of a neural progenitor to differentiation of diverse neural subtypes which restore a fully functional brain during whole-body regeneration.

## Results

### Chromosome-scale assembly and chromatin profiling reveals transcription factor binding dynamics and motifs putatively important in anterior regeneration

To conduct a comprehensive study of dynamic regulation of transcription factor binding during regeneration, we improved the genome assembly for *Hofstenia* by applying Hi-C data to achieve chromosome-scale assembly^71,72^ (Fig. S1a). The assembly totaled 992 Mb of sequence assembled in eight chromosomes (Supplementary table 1). Next, we annotated this genome sequence with a curated list of transcription factor binding motifs (Supplementary table 2). This list collapsed potentially redundant motifs that correspond to vertebrate-specific duplications of transcription factor families to consensus motifs. For example, the NK homeobox motifs corresponding to Nkx1, Nkx2, and Nkx4 were represented as one motif for the entire Nkx family in *Hofstenia*. This curation resulted in a list of 78 non-redundant motifs for distinct transcription factor protein families.

Next, we utilized the assay for transposase-accessible chromatin with sequencing (ATAC-seq) data to study the dynamic patterns of transcription factor binding at these motifs over the course of regeneration in both head and tail fragments (Fig. 1b). We mapped previously-published ATAC-seq datasets from head and tail regenerating wound sites at “0” hours post amputation (hpa), to capture chromatin state at the moment of amputation, and 6hpa^55^ as well as a newly-generated 1hpa timepoint. We identified regions of chromatin that were significantly open relative to background using Genrich. Within these regions, we used TOBIAS^73^ to calculate the binding score, because occupancy of motifs by TFs results in a specific pattern of read mapping that is detectable in our ATAC-seq data. We noted that HES1, CAMTA1, XBP1, bZIP, and NFYA motifs showed the highest levels of binding at a genome-wide scale across all time points. Among these, the motif for the Nuclear Factor Y (NFY) family of TFs stood out having the highest binding scores (Fig. 1b). Notably, despite having high binding scores relative to other motifs at all timepoints, NFY motifs appeared to be dynamically bound, becoming substantially more bound 1hpa upon amputation relative to 0hpa, the control sample (Fig. 1c). This dynamic binding was detectable specifically in tissue from tail fragments that would regenerate new heads, and this asymmetry was observable until 12hpa (Fig. S1c-d). In contrast, dynamic binding in the regenerating head fragments at 1hpa relative to 0hpa revealed that the JUN motif is differentially bound (Fig. S1b). Corroborating the previously-studied role of Egr in both head and tail regeneration in *Hofstenia*, the EGR motif was differentially bound in both regenerating head and tail fragments at 3hpa and 12hpa relative to 0hpa (Fig. S1c-d).

Given their high degree of binding relative to other motifs, we focused on NFY motifs. To uncover the biological processes that might be under the control of NFY binding sites, we applied Gene Ontology Enrichment analysis to the top 10% genes that show the largest changes in binding, going from low to high binding scores between 0hpa and 1hpa (Supplemental table 3). We found that these genes were enriched for neuronal functions (Fig. 1d). This is consistent with the dynamism of these sites being specific to tail fragments making new heads – since the brain is concentrated within the anterior, tail fragments must produce substantially more neuronal tissue relative to head fragments, which do not need to regenerate a new brain. Our observations are consistent with the hypothesis that one major role for the NFY family of transcription factors could be to control neuronal regeneration, and therefore we next studied the functions of these factors during whole-body regeneration.

### *NFY* is required for regeneration of anterior structures, including the brain

To investigate the function of *NFY* during whole-body regeneration, we assessed orthology of the NFY complex components (Fig. S2a, Supplementary Table 4), determined the spatial and temporal patterns of NFY component expression (Fig. S2b), and performed *NFY* RNAi (Fig. 2a-c, Fig. 2d-f). Using fluorescent *in situ* hybridization (FISH), we visualized mRNA expression of the members of the NFY complex (*NFYA*, *NFYB1*, *NFYB2*, and *NFYC*) (Fig. S2b-c). *NFYA* showed wound-induced expression in regenerating head and tail fragments at 6hpa (Fig. S2b). *NFYB1* and *NFYC* show wound induced expression in regenerating head and tail fragments at 15hpa and *NFYB2* shows wound induced expression in tail fragments at 6hpa and then in both head and tail fragments at 15hpa (Fig. S2c). In the NFY RNAi condition, we combined all members of the NFY complex (*NFYA*, *NFYB1*, *NFYB2*, and *NFYC*), which in other metazoans form a complex where NFYA binds DNA^74^, together to ensure complete knockdown. We performed RNAi and noticed that *NFY* RNAi regenerating tail fragments made a blastema 7dpa (Fig. 2a, Fig. S2d) with a small number of fragments showing a visible mouth (Fig. S2d), albeit at a low number. One striking phenotype we identified in the regenerating *NFY* RNAi tail fragment was their aberrant swimming behavior and overall locomotion (Fig. 2b, Supplementary Movies 1a-b, Supplementary Table 5). *NFY* RNAi regenerating tail fragments exhibited atypical locomotion, including inchworm-like movements where animals would start-and-stop, have contractions, and showed an overall decrease in swimming behavior, reminiscent of neural phenotypes shown in planaria^40,42,44,75–80^. With the aberrant swimming behavior and gene ontology analysis (Fig. 1d) implicating an issue with regenerating the nervous system, we assessed the expression of genes found with concentrated expression in the brain, *gad-1* and *TrpC-1*, using FISH. *gad-1* and *TrpC-1* are well documented in the *Hofstenia* nervous system^53^ and label neurons concentrated in the brain as well as in the body. Previous work on the neural architecture of *Hofstenia* shows a detailed timeline of neural regeneration where both *gad-1* and *TrpC-1* expression and major neural structures return by 7dpa^53^. Strikingly, *NFY* RNAi regenerating tail fragments failed to recapitulate a structure known as the anterior condensation, also referred to as the brain^53–55^ by 7dpa (Fig. 2c).

**Figure 2.**
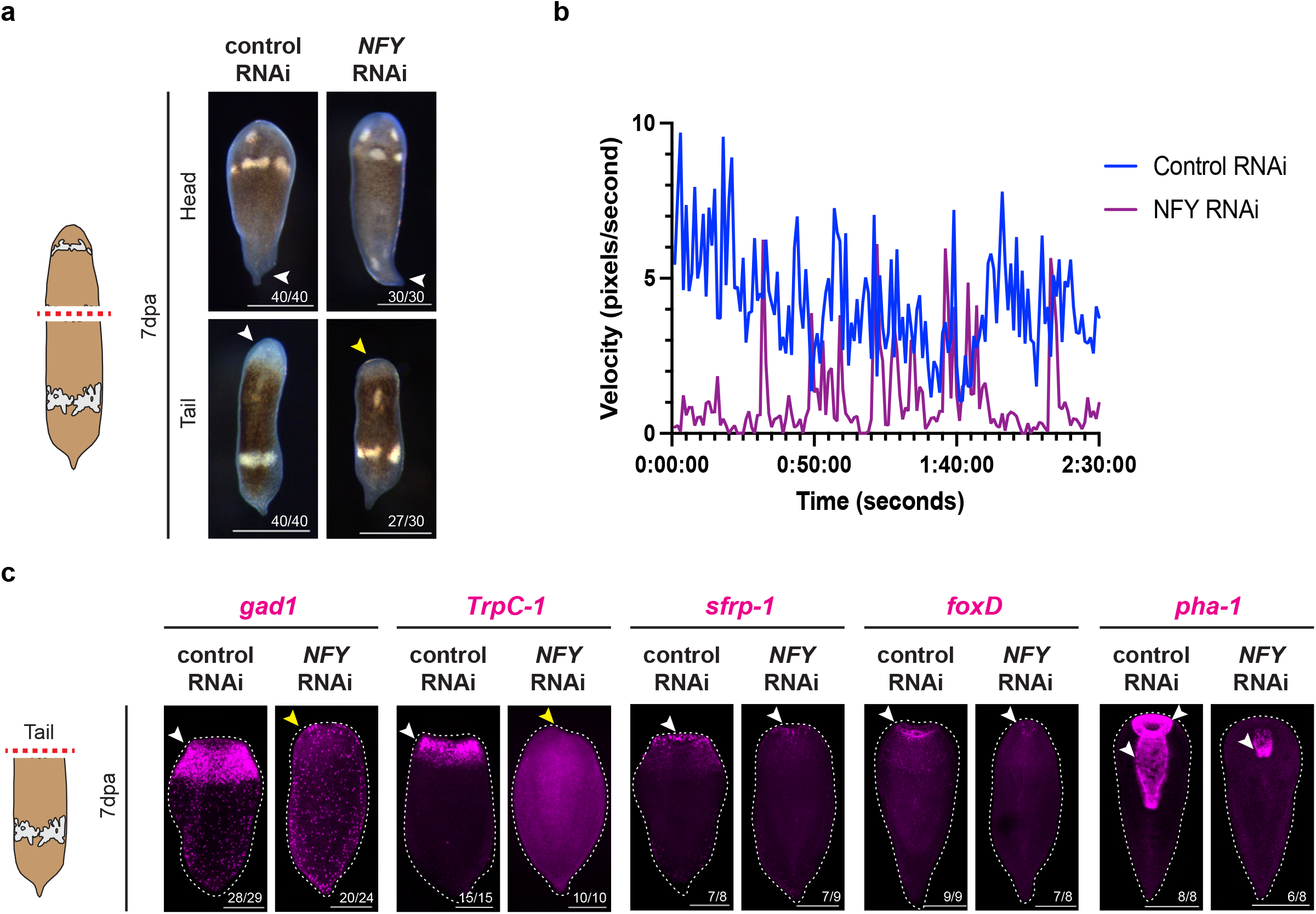
*NFY* RNAi drastically impacts the nervous system regeneration, with some other anterior structures also impacted. (A) *NFY* (consisting of *NFYA*, *NFYB1*, *NFYB2*, and *NFYC*) RNAi animals make a blastema. Regenerating head and tail fragments are shown 7 days post amputation (7dpa). White arrowhead indicates proper blastema in control RNAi. Yellow arrowhead indicates an atypical blastema. Scale bars 200 µm. (B) Plot showing the velocity (in pixels) of representative control regenerating RNAi animal (blue) and *NFY* RNAi regenerating animal (purple) over the course of two minutes and thirty seconds. The NFY RNAi regenerating animal shows repeated stopping behavior and an overall decreased velocity. (C) *NFY* RNAi regenerating tail fragments lack concentrated expression of neural markers *gad-1* and *TrpC-1* in the anterior but do show expression of anterior markers *sFRP-1* and *FoxD* as well the pharynx marker, *pha-1*. Regenerating head and tail fragments are shown 7dpa. These anterior and pharyngeal markers do seem impacted but not as drastically as the nervous system. White arrowheads indicate normal gene expression in control RNAi. Yellow arrowheads indicate lack of anterior neural expression. Scale bars 200 µm. Associated head fragments are in Supplementary Figure 2f.

This lack of anterior neural expression in *NFY* RNAi regenerating tails could be due to either a brain-specific phenotype or due to a more general problem with regeneration of anterior structures in tail fragments. We assessed whether a proper anterior was patterned in the regenerating tail fragments by looking at the anterior polarity markers *sfrp-1* and *foxD*^64,70^. Both *sfrp-1* and *foxD* expression returned in the anterior of *NFY* RNAi regenerating tail fragments by 7dpa, highlighting that a proper anterior has been patterned (Fig. 2c); however, these cellular populations were impacted relative to control animals, albeit not as drastically as the cellular populations associated with the nervous system. To further investigate whether another anterior structure was properly regenerated, we assessed the expression of *pha-1*, a known pharyngeal and mouth marker in *Hofstenia*^81^. In the *NFY* RNAi regenerating tail fragments, we identified expression of *pha-1*; however, it was truncated and a mouth was not present in most animals (Fig. 2c). Similarly, another pharyngeal and mouth marker, *desmoglein*, did not return in *NFY* RNAi regenerating tail fragments, supporting the idea that some animals do not properly regenerate a mouth (Fig. 2b). Expression of *desmoglein* was not impacted in *NFY* RNAi regenerating head fragments (Fig. S2e). The regenerating head fragments associated with Figure 2c were not impacted in the *NFY* RNAi condition, exhibiting proper expression of neural markers and anterior polarity (Fig. S2f). With *NFY* RNAi drastically impacting the nervous system during regeneration, we next sought to identify the major neural subpopulations present in the *Hofstenia* nervous system during whole-body regeneration and determine if they are governed by NFY.

### Identification of neuronal subpopulations, including a *sox4*^+^ neural-specialized stem cell

To identify major neural subpopulations present during regeneration, which are potentially impacted during *NFY* RNAi, we reanalyzed single-cell RNA-sequencing (scRNA-seq) data over the course of regeneration in *Hofstenia*^54^. To reconstruct neuronal regeneration, we extracted from this dataset differentiated neurons, putative neural progenitors, and neoblasts, the adult pluripotent stem cells of *Hofstenia*. All newly made cells during regeneration in *Hofstenia* are derived from neoblasts, and our objective in including putative progenitor populations was to capture cells that are on a path toward differentiating into neurons. By reclustering these cells, we recovered 20 subpopulations (Fig. 3a-b, Supplementary table 6).

**Figure 3.**
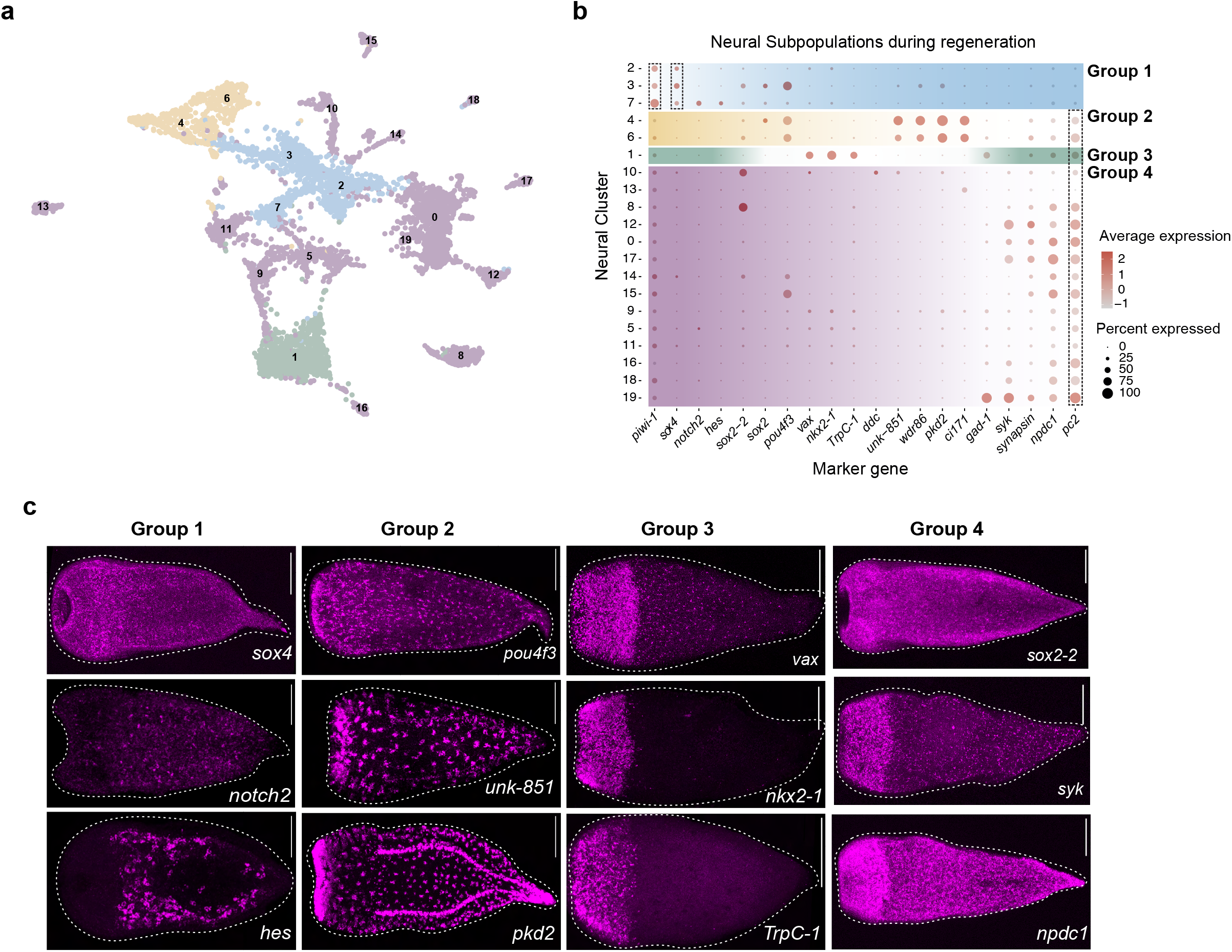
Identification of major neural subpopulations during regeneration. (A) Uniform Manifold Approximation and Projection (UMAP) representation of neural cells from a merged regeneration scRNA-seq dataset^54^ that were reclustered to identify neural subpopulations. There are 20 putative neural subtypes identified. (B) A dot plot showing the neural subpopulations on the y-axis from the UMAP with dots of varying size (the number of cells expressing a given gene) and color (the average expression of a given gene in a given cluster). The genes displayed along the x-axis represent the neoblast marker *piwi-1* and a top differentially neural gene from each neural subpopulation. There are four major neural types, falling into groups 1-4. A major transcriptional distinction between the populations is the presence of relatively high *piwi-1* expression (dotted black box on the left) and relatively high *pc2* expression (dotted black box on the right). *pc2* is also supported as a pan-neural marker because of its presence across groups 2-4. Group 1 likely represents a neural progenitor population and is the only group that expresses relatively high levels of *sox4* (dotted black box, second from the left). (C) FISH corroboration of selected markers corresponding to each of neural groups. Scale bars 200 µm. Additional FISH in Supplementary Figure 3a.

To evaluate the neural subpopulations identified here, we generated a dot plot to visualize relative expression levels of the top marker gene for each of the twenty populations as well as the neoblast marker *piwi-1* (Fig. 3b). This showed that the 20 subpopulations could be grouped into two major classes: (i) those with high-expression of the neoblast marker *piwi-1* or (ii) those with high-expression of the pan neural marker *pc2*^53^. Previous work in *Hofstenia* has shown that cells expressing relatively high levels of *piwi-1* are often associated with either neoblasts or specialized progenitor populations^54^ and that relatively high levels *pc2* expression is often associated with terminally differentiated neural cell types^53,54,64^. Neural subpopulations with high levels of *piwi-1* expression, which we refer to as Group 1, also showed relatively high expression of transcription factors (*sox4*, *notch2*, *hes1*, *sox2-2*, *sox2*, and *pou4f3*) associated with neurogenesis in other metazoan systems^15,36,42,82–86^. Subpopulations with high *pc2* expression could be further subdivided into three classes, Groups 2-4, based on distinct expression of suites of marker genes.

To further discern the identities of Group 1-4 neural subpopulations, we performed *FISH* on the top differentially expressed genes found within these groups (Fig. 3c and Fig. S3a). We found that the mRNA expression patterns for genes found within the same neural groupings showed strong correspondence. Group 1 marker genes (*sox4*, *hes*, and *notch*) labeled cells matching the distribution of neoblasts. Group 2 gene expression was found in punctate or clustered expression patterns across the body and based on gene expression associated with this group (e.g. *pou4f3*, *pkd2*, *ci171*) associated with sensory ciliated cell types as well as the patterns observed, we hypothesize these populations represent putative sensory neural subtypes. Group 3 genes (*vax*, *nkx2-1*, *TrpC-1*) labeled a prominent domain of expression in the anterior, where the brain is located. Group 4 genes, included genes for neuronal function such as *gad-1* and *synapsin*, and labeled the brain as well as cells broadly distributed the body, corresponding to the nerve net. Altogether, Groups 2-4 appear to be comprised of neural subtypes, based on expression of suites of genes consistent with terminal differentiation.

To further characterize the putative neural specialized neoblasts and neural subtypes from Groups 1-4, we computationally predicted the cell-cycle stage by using cell-cycle-associated genes^87^. We found that the subpopulations in groups 2-4 have expression associated with being in G1, or cells that are not actively cycling. Interestingly, the subpopulations found in Group 1 primarily consist of cells that have expression associated the S, G2, or M, or cells that likely about to or undergo mitosis (Fig. S3b). The subpopulations associated with Group1 have (i) relatively high levels of *piwi-1* expression, (ii) lack relatively high levels of *pc2* expression, (iii) neoblast-like expression patterns, and (iv) express genes associated with cells that are mitotically active. Together, these data led us to hypothesize that Group 1 cells represent neural specialized neoblasts. These cells would be required for the regeneration of the brain and could be impacted during *NFY* RNAi. Notably, all subpopulations in Group 1 were unified in their expression of *sox4*, suggesting that *sox4* is a marker of neural progenitors (Fig. 3b).

### *sox4* is required for the regeneration of distinct neural cell types

Given our objective of connecting the early binding dynamics of NFY to the earliest stages of neural regeneration, we examined the expression dynamics of the transcription factors associated with the *sox4^+^* neural progenitor population upon amputation. We determined that *sox4* exhibits wound-induced gene expression as early as 12hpa in regenerating tail fragments (Fig. 4a, Fig. S4b) whereas other neural transcription factors found within Group 1 (*hes*, *sox2-2*, *sox2*, and *pou4f3*) do not (Fig. S4a). We assessed *sox4* expression during a time course of regeneration and saw that *sox4* appeared to be wound induced in regenerating tail fragments at 12hpa (Fig. 4c) and that *sox4* expression in regenerating head fragments returned to a wild-type like pattern by 5dpa while regenerating tail fragments have enriched *sox4* expression in the anterior up to 7dpa (Fig. S4b). To further investigate the relationship between the putative neural progenitor population and *sox4* gene expression, we projected *sox4* in the neural subpopulations (Fig. 4b) and determined that *sox4* expression was enriched in the central UMAP clusters. We also projected *sox4* in the merged regeneration scRNA-seq data (Fig. S4c) and determined that *sox4* was expressed in cells found in the neoblast population, an intermediate population, and within the differentiated neural populations (Fig. S4b). This raised the possibility that *sox4* could be labeling a progenitor population.

**Figure 4.**
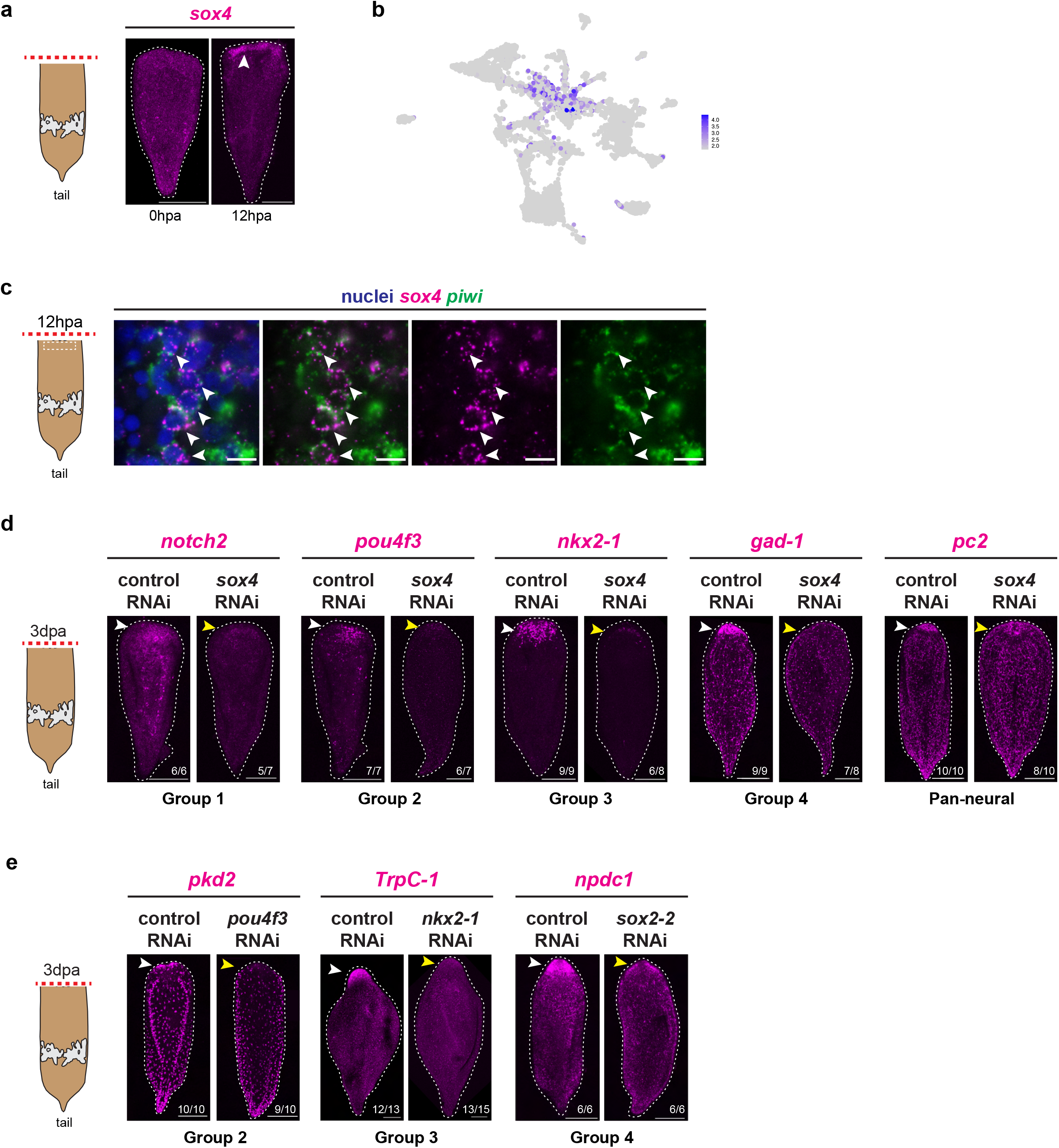
*sox4* is wound-induced in a neoblast subset and is required for the expression of neural genes associated with distinct neural subpopulation. (A) *sox4* expression is wound-induced in regenerating tail fragments at 12hpa relative to a 0hpa. White arrowhead indicated wound-induced gene expression. Scale bars 200 µm. (B) UMAP representation of neural subpopulations depicting *sox4* expression. (C) Double FISH of the putative neural progenitor marker *sox4* and neoblast marker *piwi-1* in regenerating tail fragments at 12hpa. Dotted white box indicated region of interested where high-magnification imaging was performed. Co-expression of *sox4* and *piwi-1* (denoted by white arrowheads) was detected in a subset of *piwi-1*^+^ cells. Scale bars 10 µm. (D) RNAi of *sox4* impacts gene expression associated with neural groups 1-4 and the pan-neural marker *pc2*. Regenerating tail fragments are shown 3dpa. White arrowheads indicate normal gene expression in control RNAi. Yellow arrowheads indicate impacted neural gene expression. Scale bars 200 µm. (E) RNAi of differentially-expressed putative neural transcription factors expressed within different neural groups. Regenerating tail fragments are shown 3dpa. For each group, we performed RNAi of a transcription factor associated with that group and assessed expression of a putative differentiated gene associated with the same group. White arrowheads indicate normal gene expression in control RNAi. Yellow arrowheads indicate impacted neural gene expression. Scale bars 200 µm.

Double FISH for *sox4* and the neoblast marker *piwi-1* corroborated the single-cell RNA-seq data, showing the *sox4*^+^ cells were also *piwi-1*^+^ in intact and regenerating animals. We found that in intact animals, *sox4* and *piwi-1* double-positive cells are detected, with *sox4*^+^ cells being a subset of *piwi-1*^+^ cells (Fig. S4d). In regenerating animals, we found that similar to intact animals, there are *sox4* and *piwi-1* double-positive cells with *sox4*^+^ cells being a subset of *piwi-1*^+^ cells (Fig. 4c). However, unlike expression in intact animals, there was *sox4* wound-induced expression at the wound site, largely within the neoblast compartment, as early as 12hpa (Fig. 4b). This is consistent with either proliferation of *sox4^+^* neoblasts or the activation of *sox4* expression in *piwi-1^+^* neoblasts that were previously *sox4^-^*. Mitotic activity is concentrated at the wound site ∼48-72hpa, but is not normally enriched at the wound by 12hpa^64^. Therefore, we infer that wound-induced *sox4* reflects an increase in *sox4* expression within the neoblast population at the wound site. These data provide evidence supporting that wound-induced *sox4* expression occurs in neoblasts, possibly resulting in the formation of neural progenitors.

To determine the function of *sox4* during whole-body regeneration, we performed *sox4* RNAi and assessed neural gene expression 3dpa, a time point when differentiated neurons are known to return during regeneration^53,54^. We found that *sox4* is required for the expression of marker genes for Group 1-4 neural subtypes during regeneration (Fig. 4d and Fig. S4e). Specifically, *sox4* is required for *notch2* (group 1), *pou4f3* (group 2), *nkx2-1* (group 3), *gad-1* (group 4), and *pc2* (pan-neural) expression during regeneration. Thus, genes regulated by *sox4* included transcription factors as well as genes associated with differentiated neural cell types in neural subtype Groups 1-4.

We noted that neural subtypes in Groups 2-4, which expression of *pc2* and other genes of differentiated neurons, also showed expression of different transcription factors. To determine if these transcription factors are required for the regeneration of their corresponding neural subtypes (e.g. group 2 neural transcription factor is required for group 2 differentiated neural gene expression), we performed RNAi of a transcription factor associated with a given subtype and assessed expression of a differentiated neural gene associated with it (Fig. 4e, Fig. S4e). We assessed Group 2, the putative sensory cell types, by performing RNAi of *pou4f3* and determined that it was required for *pkd2* expression during regeneration. Interestingly, the transcriptional regulation of *pkd2* by *pou4f3* has been identified in planarians^40,88^ as well as other metazoans^89–95^. For Group 3, cell types associated with the brain, we found that *nkx2-1* was required for *TrpC-1* expression during regeneration. Finally, we assessed Group 4, cell types associated with the brain and broad body, by performing RNAi of *sox2-2* and determined that *npdc1* and gad-1 expression were impacted (Fig. 4e, Fig. S4f). Together these data suggest that *sox4* is upregulated in a subset of neoblasts that are specialized to the neural lineage, and *sox4* is required for the expression of transcription factors that are further needed to generate distinct differentiated neural cells.

### NFY regulates *sox4* wound-induced expression during regeneration

Given that *sox4* was the earliest wound-induced gene needed for neural regeneration, we next sought the transcriptional regulatory decision-making that controls the *sox4* locus. We identified the ∼12.1kb region of the *Hofstenia* genome that encompasses the *sox4* gene and visualized chromatin accessibility data from regenerating tail fragments at 0, 1, and 6 hpa (Fig. 5a). We identified consensus peak sets and used DiffBind^96,97^-called regeneration peaks to determine that there is one statistically significant, differentially accessible region within the *sox4* locus when comparing data from 0hpa to 1hpa (*p* = 1.63×10^-10^). Focusing on this peak, we found an underlying NFY transcription factor binding site. Based on the TOBIAS^73^ footprinting analysis, this NFY binding site showed increased occupancy by a protein likely binding to it, with the site going from being “unbound” at 0hpa to “bound” at 1hpa (Fig. 5a). Data from 6hpa animals showed that while the accessibility of the chromatin decreases, the site likely remains bound (or in other words, occupied by a DNA-binding protein). We hypothesized that NFYA subunit of the NFY complex is likely binding at this NFY motif in the *sox4* locus at 1hpa. This one site would be one of the many NFY sites that become more bound between 0hpa and 1hpa genome-wide (heatmap Fig. 1b). These data raised the possibility that NFY could be a direct transcriptional regulator of *sox4* expression, such that the dynamic binding of the NFY motif in the *sox4* locus leads to wound-induced *sox4* expression.

**Figure 5.**
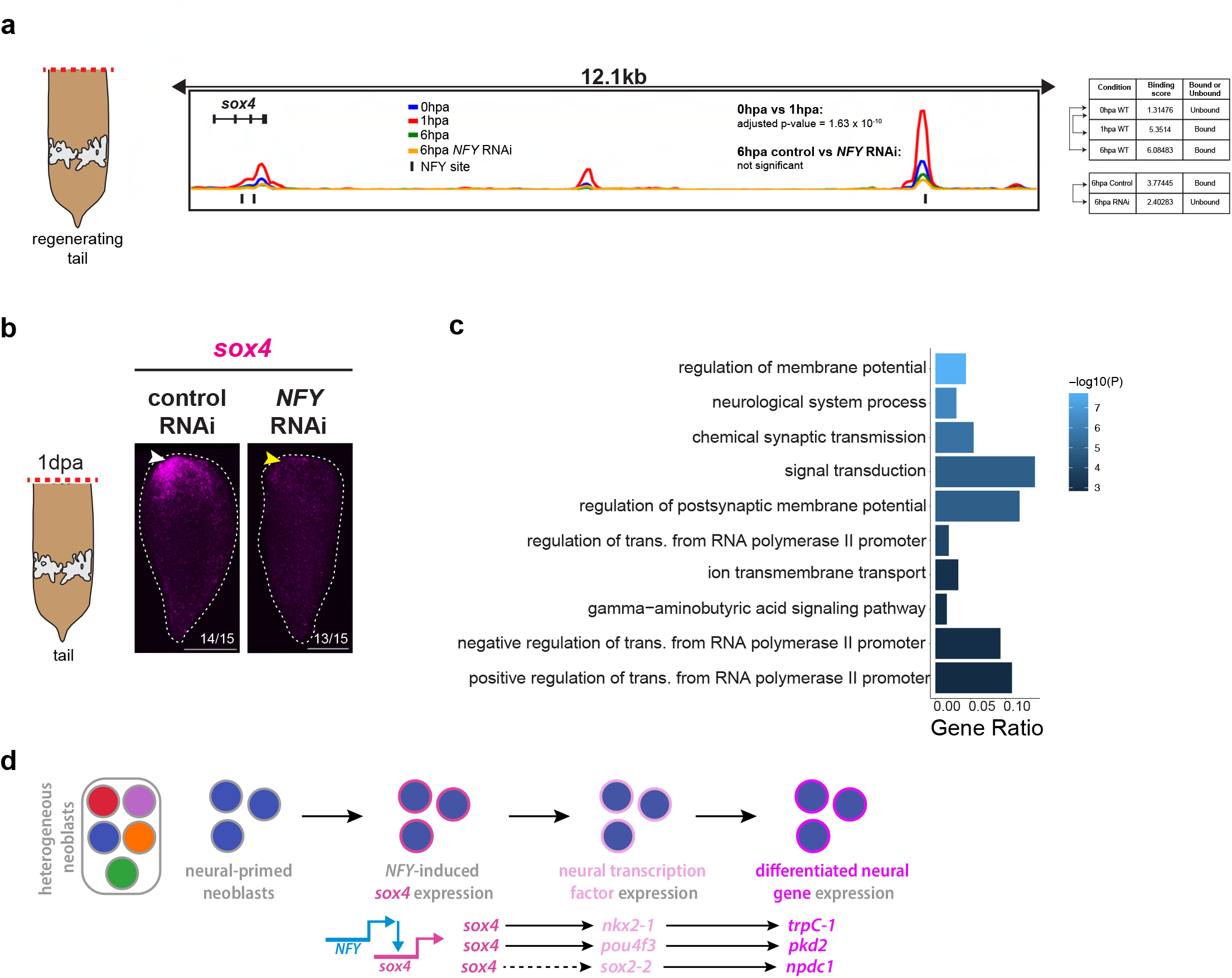
NFY wound-induced transcription factor binding regulates *sox4* expression during regeneration. (A) Left: Schematic depicting the ∼12.1kb region that encompasses the *sox4* gene locus showing ATAC-seq data for 0, 1, and 6 hour regenerating tail fragments along with sites that are putatively bound by the NFY transcription factor (at the NFYA site). The NFYA site on the right is of interest because it is initially unbound at 0hpa and becomes statistically significantly bound at 1hpa. Right: Table of conditions, binding scores, and whether the specific NFYA site is bound or unbound. Arrows along the left side indicate which samples were compared using TOBIAS to assess relative binding. (B) RNAi of *NFY* leads to a lack of *sox4* upregulation in the anterior of regenerating tail fragments 1dpa. White arrowhead indicates *sox4* upregulation in control RNAi. Yellow arrowhead indicates a failure of *sox4* upregulation. Scale bars 200 µm. (C) In the *NFY* RNAi ATAC-seq data, dynamically bound NFY sites located proximal to genes associated with neural function are no longer accessible at 6hpa. Gene ratio is the number of genes in a particular biological process category over the total list of genes. (D) A proposed model of NFY wound-induced transcription factor binding which leads to the upregulation of *sox4* gene expression in neoblasts during regeneration which is required for the expression of genes associated with distinct neural subpopulations.

To corroborate that *NFY* is regulating *sox4* wound-induced expression during regeneration, we performed RNAi of *NFY* and assessed expression of *sox4*. We determined that *sox4* fails to be upregulated in regenerating tail fragments assessed at 1dpa upon *NFY* RNAi (Fig. 5b). To assess if transcription factor binding at the NFY site in the *sox4* locus is impacted under *NFY* RNAi, and to identify more targets of NFY genome-wide, we obtained ATAC-seq data from *NFY* RNAi and control RNAi animals at 6hpa. First, using TOBIAS^73^ footprinting analysis of the 6hpa control RNAi (binding score = 3.77445) versus the 6hpa *NFY* RNAi (binding score = 2.40283), we determined that in the *NFY* RNAi condition, the NFY motif is unbound at 6hpa relative to control RNAi (Fig. 5a, Supplementary Table 7). Second, *NFY* RNAi ATAC-seq data yielded binding scores for all NFY binding sites in the genome under NFY and control RNAi. We focused on genes associated with the top 10% of *NFY* sites that showed diminished binding in the *NFY* RNAi condition compared to the control RNAi condition at 6hpa, as a conservative list of putative NFY target genes. Gene ontology enrichment analysis of these genes determined that NFY targets are enriched for functions in the nervous system (Fig. 5c). Thus, in addition to directly regulating *sox4*, NFY appears to mediate direct transcriptional control over many neural genes.

## Discussion

Here, our work identifies a path from wounding to activation of a neural progenitor to differentiation of diverse neural subtypes that restore a fully functional brain during whole-body regeneration in an adult animal. Specifically, our results suggest a model in which wound-induced binding of NFY to NFY transcription factor binding sites in the *Hofstenia* genome occurs as early as 1hpa with an enrichment of these sites located in close proximity to neural related genes. NFY then induces *sox4* expression (likely in neoblasts) which is required for the regeneration of transcriptionally distinct neural subpopulations (Fig. 5d). Work in diverse metazoan systems focusing on neurogenesis during whole-body regeneration has focused on identification of differentiation pathways. Studies in the planarian *S. mediterranea* identified putative neural lineages using *in silico* trajectory inference, including a neoblast subset associated with the nervous system^98^. In *Hydra*, putative differentiation trajectories within the nervous system have been identified highlighting similarities between neurogenesis and gland cell differentiation suggesting a shared or similar progenitor state^22^. Our work in the acoel *Hofstenia miamia* reveals a wound-induced transcription regulatory mechanism, where NFY becomes bound to its motifs within an hour of amputation, and ultimately facilitates the specification of a *sox4^+^* neural stem cell population, as a necessary step in a wounding-induced pathway for neurogenesis.

NFY has many roles in other metazoan systems including fate specification by binding to cell type specific enhancers along with cell-type specific transcription factors^99^. NFY has been extensively studied with regard to cell cycle progression and proliferation through the binding and direct activation of target genes^100^. In a variety of vertebrate systems, the NFY complex is required for maintaining stem cell identities with the hematopoietic stem cell system^101^, in skeletal muscle stem cells^100,102^, as well as caridiomyocytes^103^. In planarians, NFY is required for the self-renewal and proliferation of spermatogonial stem cells but not for their specification or differentiation^104^. We discovered a novel role for NFY during whole-body regeneration in the and for its function regarding the activation of neural-related gene expression in the acoel *Hofstenia miamia*.

Further, we identified genes required for neuronal differentiation during whole-body regeneration, including *sox4* which is required for the regeneration of diverse neural subtypes. *sox4* is a *soxC* (Sry-related HMG box transcription factor) group member known to play an important role in different cellular contexts including neuronal cell fate, especially in embryonic neural progenitors across vertebrates and invertebrates^105^ as well as a role in neuronal subtype specification^106,107^. Strikingly in echinoderms, robust lineage tracing in the larvae of *Patiria miniata* has identified that *sox4^+^* neural precursors proceed along an embryonic neurogenesis pathway to regenerate the anterior nervous system^28^. In *Nematostella*, a *SoxC* ortholog was discovered as an upstream regulator of a neural progenitor population that differentiates into cnidocytes, gland cells, and neurons during development and homeostasis^52^. Intriguingly, we identified another conserved transcriptional relationship between a *pkd2^+^* neural subtype regulated by a *pou4* ortholog, a relationship identified in planarians and other metazoans where *pou4* orthologs are shown to function as regulatory genes that govern ciliated sensory neuron identity^88–95^. These seemingly conserved transcriptional relationships beg exploration and identification of the GRNs underlaying neuronal cell-type specification across more diverse animal linages.

The chromosome-scale genome assembly annotated with a curated list of binding sites and the chromatin accessibility data generated here will be powerful for obtaining mechanistic insight into other processes. Our study focused on the most strongly bound motif during whole-body regeneration, but motifs further down the list are likely to be important mediators of regeneration. The next steps will be to identify how the many wound-induced binding events connect to each other and how they initiate the differentiation of cell types that construct other organs, beyond neurons and the brain.

## Supplementary Figure Legends

**Supplementary Figure 1.**
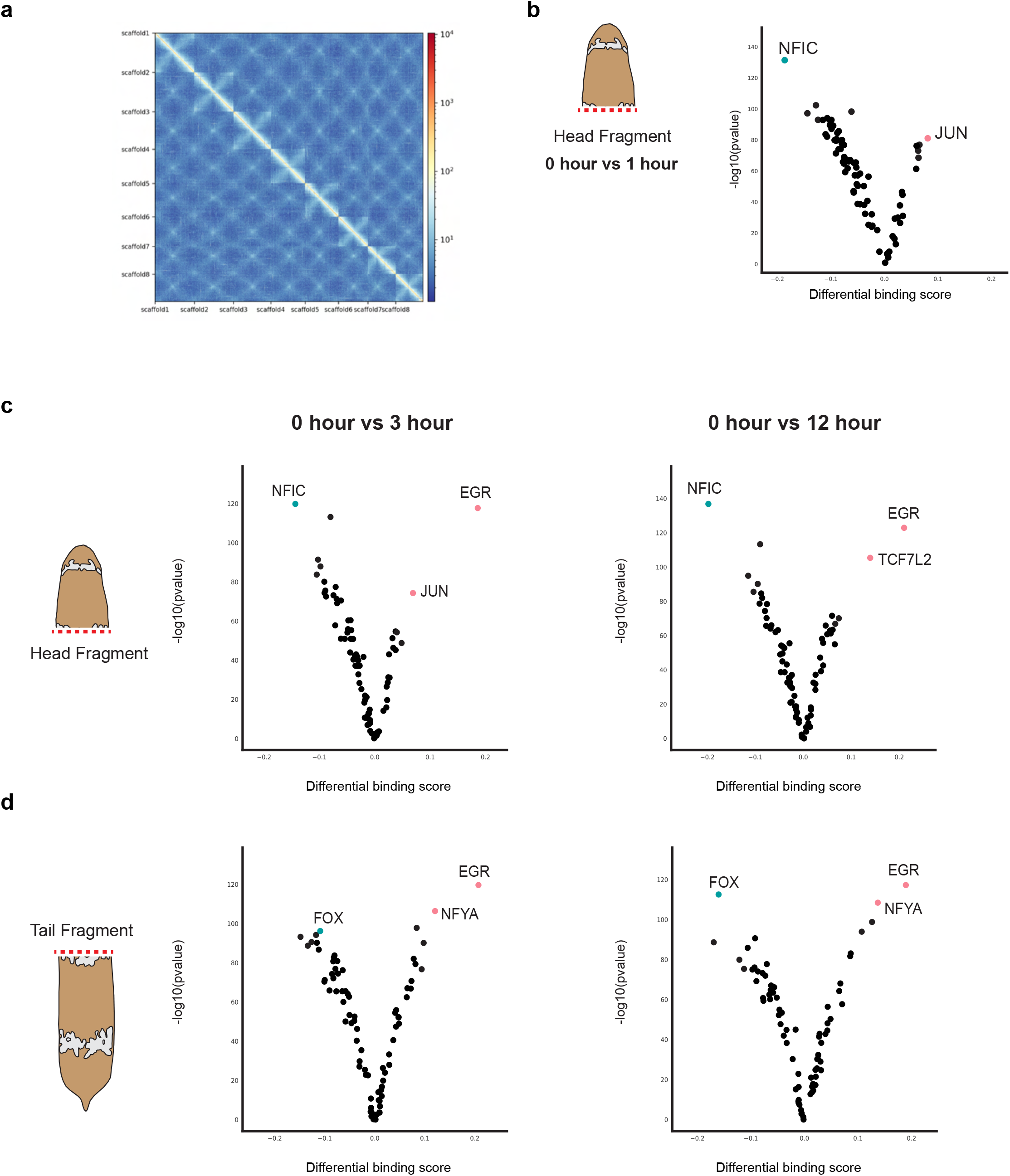
Chromosome level assembly and chromatin dynamics during regeneration. (A) Heatmap showing chromosome level genome assembly with 8 scaffolds. (B) Volcano plot showing that NFIC is the most bound motif at 0hpa and JUN is the most bound motif at 1hpa in the regenerating head fragment in the *Hofstenia* genome. (C) Volcano plot showing that NFIC is the most bound motif at 0hpa and EGR is the most bound motif at 3hpa and 12hpa in the regenerating head fragment in the *Hofstenia* genome. (D) Volcano plot showing that FOX is the most bound motif at 0hpa and EGR is the most bound motif at 3hpa and 12hpa in the regenerating tail fragment in the *Hofstenia* genome.

**Supplementary Figure 2.**
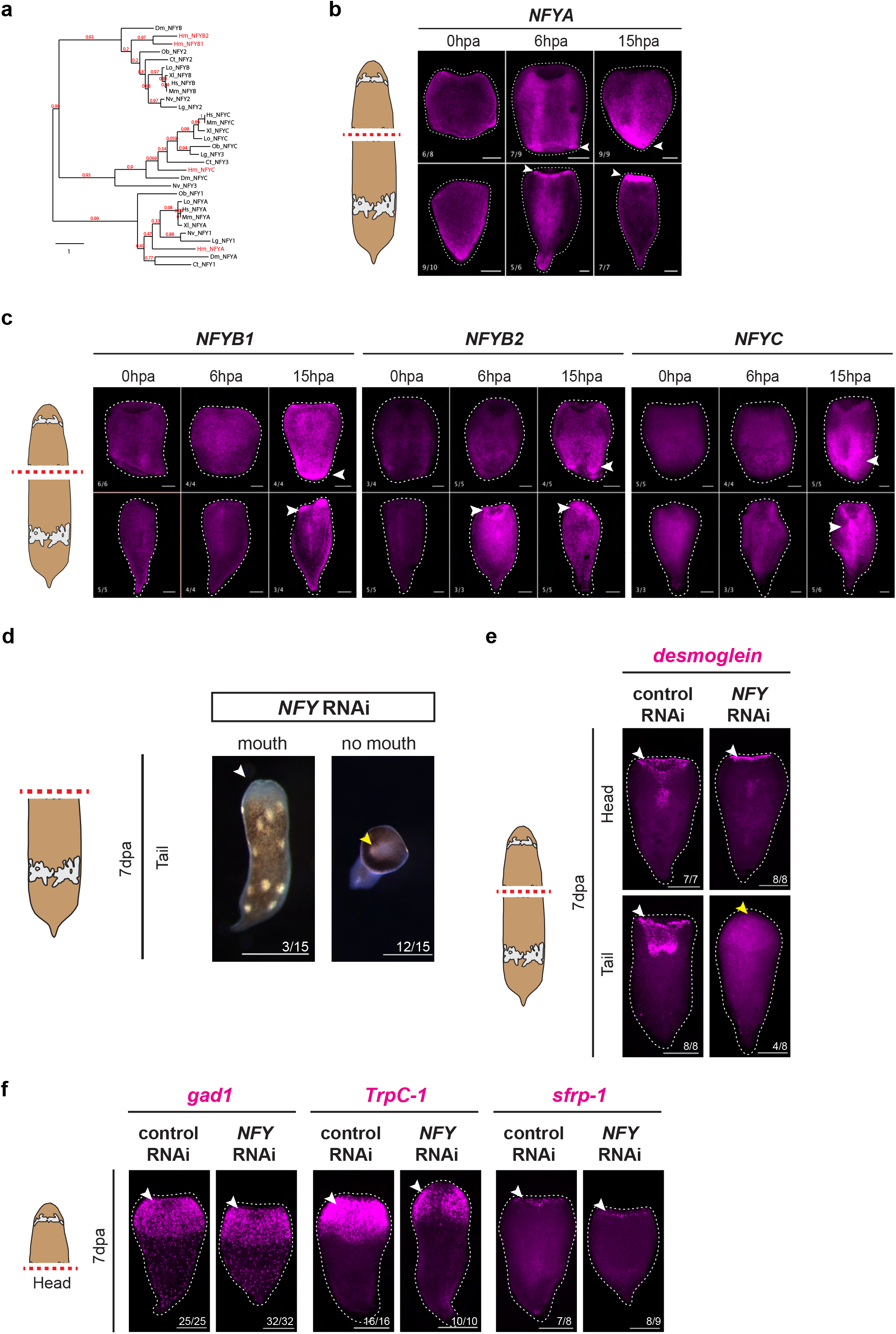
NFY complex orthology assessment, expression dynamics, and RNAi. (A) Phylogeny of the *Hofstenia* NFY complex components. (B) Regeneration time course of head and tail fragments with FISH of *NFYA* showing wound induced expression. Scale bars 200 µm. (C) Regeneration time course of head and tail fragments with FISH of *NFYB1*, *NFYB2*, *and NFYC* showing wound induced expression. Scale bars 200 µm. (D) *NFY* RNAi regenerating tail fragments 7dpa depicting low proportion of animals with a mouth (white arrowhead) and a majority of animals without a mouth (yellow arrowhead). Scale bars 200 µm. (E) *NFY* RNAi regenerating tail fragments lack expression mouth and pharyngeal marker, *desmoglein*, in the anterior. Regenerating head and tail fragments are shown 7dpa. White arrowheads indicate normal gene expression in control RNAi. Yellow arrowheads indicate lack of *desmoglein*. Scale bars 200 µm. (F) Associated RNAi head fragments are corresponding to figure 2C. *NFY* RNAi regenerating head fragments do not show an impact on the assessed markers. White arrowheads indicate normal gene expression in control RNAi. Scale bars 200 µm.

**Supplementary Figure 3.**
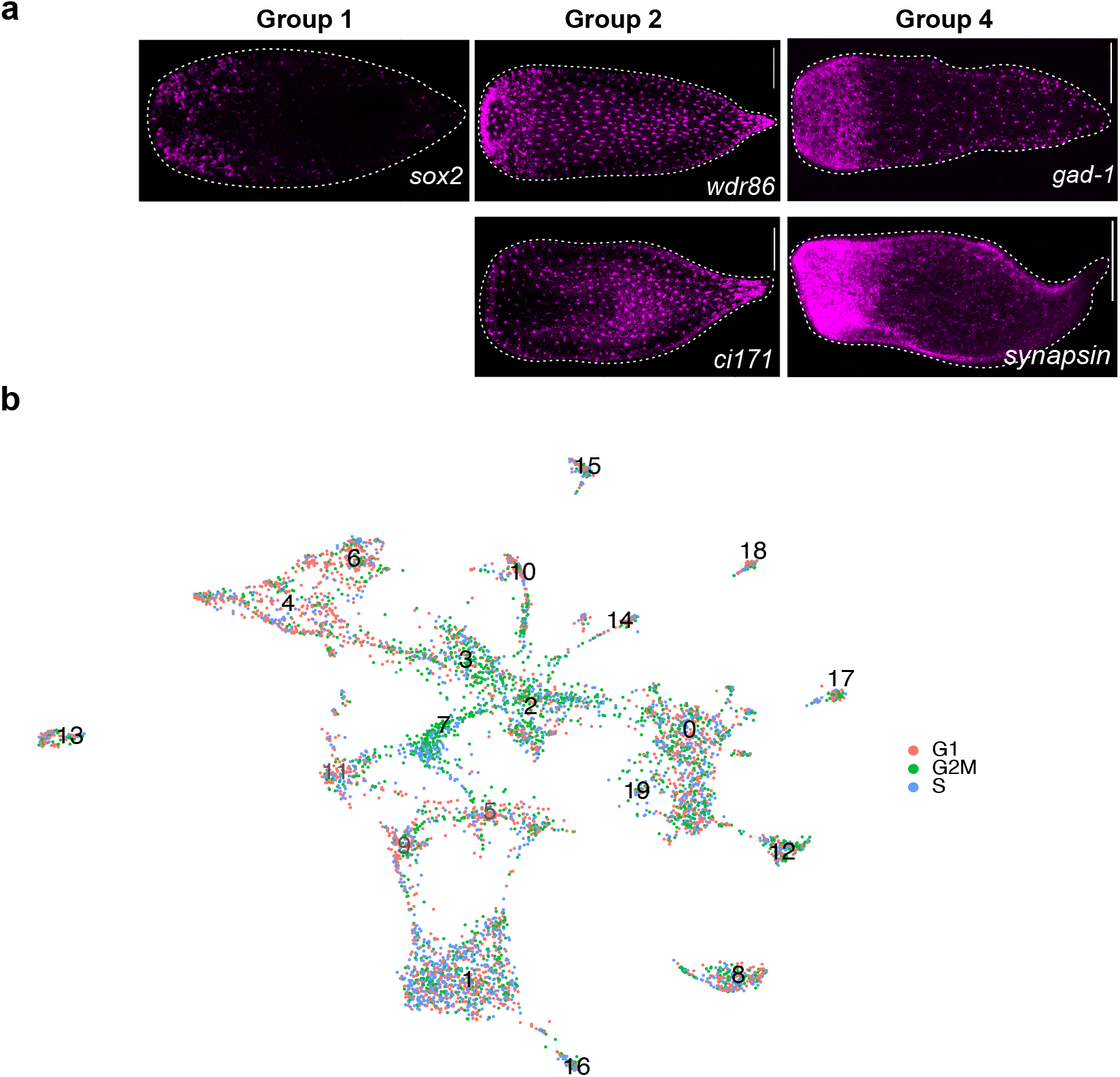
Neural subpopulation gene expression and cell-cycle scoring. (A) Additional gene expression studies for genes associated with neural subpopulations 1, 2, and 4. (B) Cell cycle scoring of neural subpopulations.

**Supplementary Figure 4.**
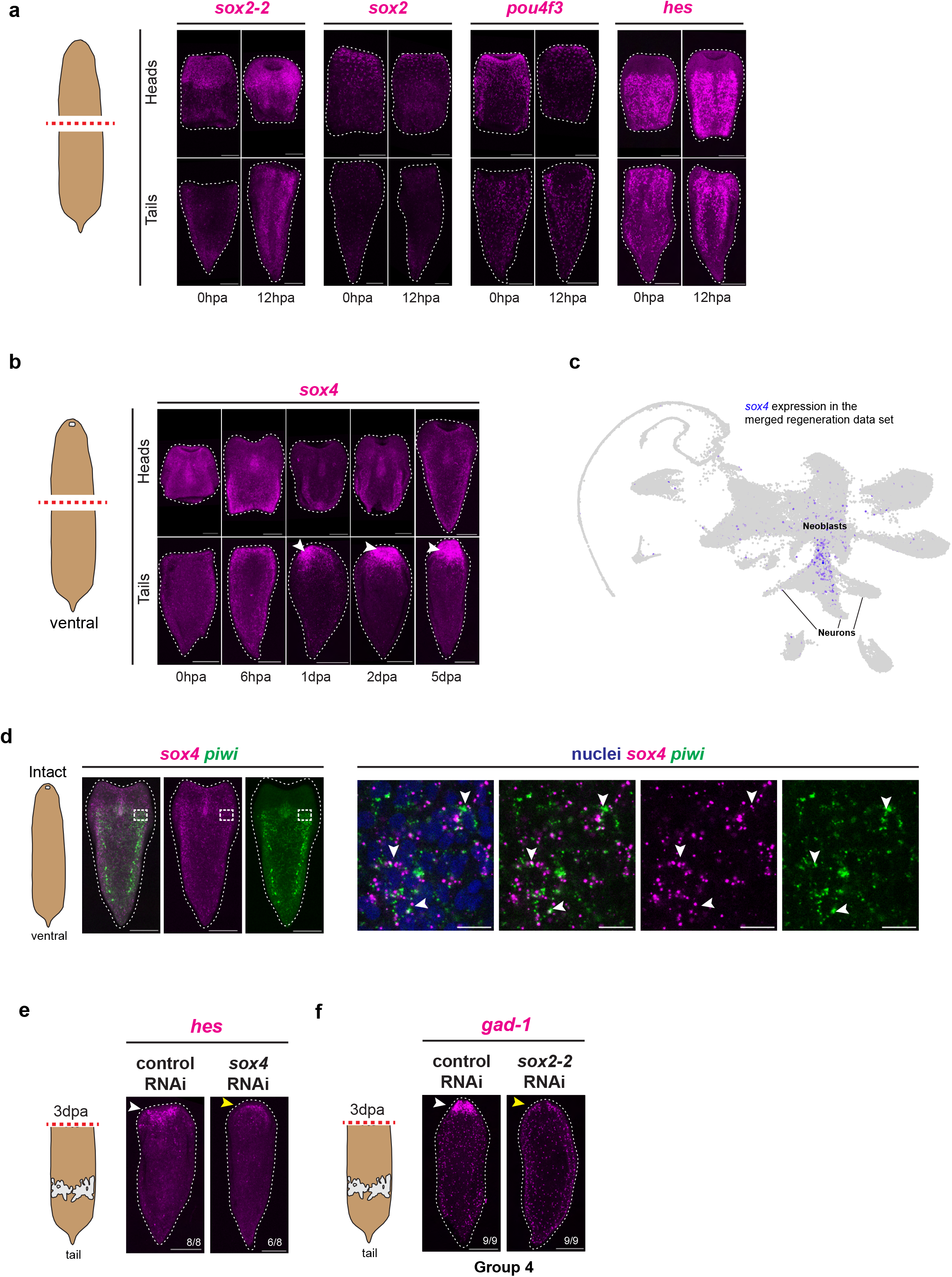
sox4 gene expression is wound-induced, found in neoblasts, and important for neural regeneration in tail fragments. (A) Group 1 transcription factors do not show wound induced expression at 12hpa relative to 0hpa in regenerating head and tail fragments. Scale bars 200 µm. (B) Regenerative time course of sox4 gene expression in head and tail fragments. White arrowheads depict enriched expression of sox4 in the anterior region of regenerating tail fragments from 1dpa to 5dpa. Scale bars 200 µm. (C) *sox4* expression projected into the merged regeneration single cell atlas. (D) Double FISH of the putative neural progenitor marker *sox4* and neoblast marker *piwi-1* in intact animals. Dotted white box indicated region of interested where high-magnification imaging was performed. Co-expression of *sox4* and *piwi-1* (denoted by white arrowheads) was detected in a subset of *piwi-1*^+^ cells. Scale bars 200 µm for entire animal. Scale bars 10 µm for zoom-ins. (E) RNAi of *sox4* impacts *hes* expression during regeneration. White arrowheads indicate normal gene expression in control RNAi. Yellow arrowheads indicate impacted neural gene expression. Scale bars 200 µm. (F) For group 4, RNAi of *sox2-2* impacts *gad-1* expression during regeneration. White arrowheads indicate normal gene expression in control RNAi. Yellow arrowheads indicate impacted neural gene expression. Scale bars 200 µm.

## Supplementary tables

Supplementary table 1: Genome assembly summary

Supplementary table 2: Consensus Transcription factor binding motifs

Supplementary table 3: TOBIAS footprinting analysis for NFY motifs in wild-type regenerating animals

Supplementary table 4: NFY orthologs and neural related genes

Supplementary table 5: Control and *NFY* RNAi swimming measurements

Supplementary table 6: Marker genes for neural subpopulations

Supplementary table 7: TOBIAS footprinting analysis for NFY motifs in RNAi animals

## Methods and materials

### Improved genome assembly

High molecular weight genomic DNA was prepared from approximately 5 hatchling sized clonal worms, which were derived from regenerated fragments of the same single founder worm as the original draft genome^55^. Genomic DNA was subsequently sequenced on an Oxford Nanopore MinION machine. Three new flowcells of data were combined with two previous runs^55^ to produce the raw long read sequencing reads, representing approximately 27X coverage. Basecalling was performed using the gpu version of Guppy (v6.0.1), assembled using Shasta (v0.1.0), and polished using previously published short reads^55^ using Pilon (v1.23). This resulted in a 992 Mb assembly with a contig N50 of 495kb; we measured BUSCO completeness using BUSCO v3.0.2 and the ‘eukaryota_odb9’ dataset, with 90.7% of genes represented in the assembly (88.1% single copy, 2.6% duplicates). To improve contiguity, we performed Hi-C using approximately 15 worms as input into the Dovetail Hi-C Library Prep kit. We performed and sequenced two replicates of Hi-C, which combined with the nanopore based assembly were sent to Dovetail for scaffolding and improved continuity. BUSCO results of the 8 largest scaffolds of the completed assembly resulted in 91.8% complete. Genome annotations were lifted from the original assembly^55^ to the new assembly using Liftoff (v0.4.3.2)

### Consensus motifs

In order to create a non-redundant list of transcription factor binding motifs, we used the genome annotation as input into Panther (v17.0)^109^ to get a list of transcription factors (TF) present in the *Hofstenia* genome. We then manually checked each TF name against the JASPAR database and assigned a position weight matrix for each putative TF. Motifs that were indistinguishable were grouped (i.e. NKx) in order to streamline motif analysis. This resulted in a list of 78 TF motifs that were used for subsequent analysis. TF binding analysis and footprinting were performed with TOBIAS (v0.11.1)^73^.

### ATAC-seq library preparation and analysis

ATAC-seq was performed as described previously, with minor modifications^55,110^. Fragments from wound sites from individual *Hofstenia* animals were isolated and placed directly into ice cold lysis buffer and homogenized manually with a pipette. This cell homogenate was filtered with a 40 µm cell strainer to obtain a single-cell solution which was then processed using a standard ATAC experimental pipeline^110^. Biological replicates were performed in duplicate (at least) for all ATAC experiments.

### Animal maintenance

Adult animals were kept in plastic boxes at 21°C in artificial seawater (38 ppt, pH 7.9–8.0; Instant Ocean Sea Salt). Juvenile worms were kept in zebrafish tanks. Seawater was replaced twice a week. Juvenile and adult worms were fed with rotifers *Brachionus plicatilis* and freshly hatched brine shrimp *Artemia sp.* twice a week, respectively.

### Phylogenetic analysis

NFY components and transcription factors of interested were identified in the *Hofstenia* genome and assigned orthology. First, a BLAST was performed to identify putative gene assignment. To assign orthology, phylogenetic trees were constructed. Sequences were aligned with MUSCLE (v3.8.31)^111^. Alignments were trimmed using Gblocks^112,113^ with the least stringent parameters Phylogenetic trees were inferred using Maximum Likelihood analysis with 1,000 bootstrap replicates, implemented in RAxML (v8.2.4)^114^ using the WAG+G model of protein evolution.

### Fixation and in situ hybridization

Whole worms and regenerating fragments were fixed in 4% paraformaldehyde in 1% phosphate-buffered saline with 0.1% Triton-X (PBST) for one hour at RT on a nutator and stored in 100% methanol at -20 °C until use. Digoxigenin and Fluorescein labeled riboprobes were synthesized as previously described ^64^. Fluorescence *in situ* hybridizations (FISH) were performed following the protocol described^64^.

### RNAi

Double-stranded RNA (dsRNA) synthesis was made following a protocol described in previous work^64^. RNAi experiments were done by injecting the animals with dsRNA corresponding to the target gene into the gut for 3 consecutive days. dsRNA injections were performed using a Drummond Nanoject II. Animals were cut transversally at least 3 hours after the third injection and were allowed to regenerate for the appropriate time while being monitored for visible phenotypes or external defects. Control dsRNA for gene inhibition was the *unc22* sequence from *C. elegans* that is absent in *Hofstenia*. Animals were fixed at different time points upon amputation and used in FISH.

### Locomotion behavior analysis

Control and *NFY* RNAi 7dpa regenerating tail fragments videos were recorded using a Leica DFC7000. We performed manual tracking of individual fragments by tracking a point at the anterior-central portion of the regenerating fragment using FIJI^115^. This point was tracked for 150 seconds and the raw data output included distance moved over time, which was plotted for the control and *NFY* RNAi fragments.

### Gene ontology enrichment analysis

Gene ontology enrichment analysis was done using the same methods as previously described^81^. To determine biological processes associated with bound NFY sites at 6hpa we usd the 6hpa regenerating tail ATAC-seq data set and analyzed the top differentially bound NFY sites in the *Hofstenia* genome. From this list of sites, we used UROPA^116^ to link these bound NFY sites to nearby genes. From this list of genes, we performed gene ontology analysis and determined that a majority of the associated biological processes are neural-related. To determine biological processes impacted during *NFY* RNAi, we used the 6hpa control and 6hpa *NFY* RNAi regenerating tail ATAC-seq data sets and analyzed the top differentially bound NFY sites in the *Hofstenia* genome in the 6hpa control compared to the 6hpa *NFY* RNAi. From this list of sites, we used UROPA to link the no longer-bound NFY sites to nearby genes and determined that a majority of the associated biological processes are neural-related.

### Single-cell RNA-seq analysis

We subclustered the neural population from a merged regeneration scRNAseq data set^54^ using a standard method from Seurat v3^117^. We determined if neural subpopulations of the UMAP were either over- or under-clustered based on known neural marker gene expression. We iteratively altered clustering parameters and UMAP rendering to make sure the results were robust and representative of the known biological cell types. Clustering was performed in an iterative manner where we identified top marker genes and projected them back onto the UMAP plots to gauge specificity and uniformity of expression within their corresponding clusters. If the top marker genes were found to be highly expressed in multiple clusters it was suggestive of over-clustering. If we found top marker genes that were only expressed within a subset of the corresponding cluster, it was suggestive of under-clustering. We also performed FISH corroboration of these findings using the top marker genes identified using the aforementioned parameters.

## Acknowledgements

We want to thank the Srivastava lab and Koenig lab for helpful discussions regarding the work. As well as the Harvard Bauer Core for their assistance in sequencing the ATAC-seq libraries. P. Grayson for help and guidance in data analysis. Y. Jyun-Luo, A. Knecht, and M. He for preliminary investigation of putative neural markers. K. Loubet-Senear for help and discussions with visualizing ATAC-seq data. D. Khost and T. Sackton for assistance with the nanopore genome assembly.

## Author Contributions

R.E.H, A.R.G., and M.S. conceived and designed the study. R.E.H. and M.S. wrote the paper. A.R.G. generated the ATAC-seq data and conducted preliminary analysis. R.E.H. performed the reanalysis and clustering of the single-cell data. C.R.L. and R.E.H. reanalyzed the *NFY* RNAi ATAC-seq data. R.E.H., A.R.G., and A.G. performed FISH corroboration. R.E.H., A.R.G., and A.G. performed confocal imaging. R.E.H. performed RNAi. A.R.G. and R.E.H. performed phylogenetic analysis of genes of interest.

